# Locus-Specific Enhancer Hubs And Architectural Loop Collisions Uncovered From Single Allele DNA Topologies

**DOI:** 10.1101/206094

**Authors:** Amin Allahyar, Carlo Vermeulen, Britta A.M. Bouwman, Peter H.L. Krijger, Marjon J.A.M. Verstegen, Geert Geeven, Melissa van Kranenburg, Mark Pieterse, Roy Straver, Judith H.I. Haarhuis, Hans Teunissen, Ivo J. Renkens, Wigard P. Kloosterman, Benjamin D. Rowland, Elzo de Wit, Jeroen de Ridder, Wouter de Laat

**Affiliations:** Department of Genetics, Center for Molecular Medicine, University Medical Center Utrecht, 3584 CG Utrecht, The Netherlands. Utrecht University, 3508 TC Utrecht; Delft Bioinformatics Lab, Faculty of Electrical Engineering, Mathematics and Computer Science, Delft University of Technology, 2628 CD Delft, The Netherlands.; Hubrecht Institute-KNAW and University Medical Center Utrecht, Uppsalalaan 8, 3584CT Utrecht, the Netherlands; Division of Gene Regulation, Netherlands Cancer Institute, Plesmanlaan 121, 1066CX Amsterdam, the Netherlands; Utrecht University, Domplein 29, 3512 JE Utrecht

## Abstract

Chromatin folding is increasingly recognized as a regulator of genomic processes such as gene activity. Chromosome conformation capture (3C) methods have been developed to unravel genome topology through the analysis of pair-wise chromatin contacts and have identified many genes and regulatory sequences that, in populations of cells, are engaged in multiple DNA interactions. However, pair-wise methods cannot discern whether contacts occur simultaneously or in competition on the individual chromosome. We present a novel 3C method, Multi-Contact 4C (MC-4C), that applies Nanopore sequencing to study multi-way DNA conformations of tens of thousands individual alleles for distinction between cooperative, random and competing interactions. MC-4C can uncover previously missed structures in sub-populations of cells. It reveals unanticipated cooperative clustering between regulatory chromatin loops, anchored by enhancers and gene promoters, and CTCF and cohesin-bound architectural loops. For example, we show that the constituents of the active b-globin super-enhancer cooperatively form an enhancer hub that can host two genes at a time. We also find cooperative interactions between further dispersed regulatory sequences of the active proto-cadherin locus. When applied to CTCF-bound domain boundaries, we find evidence that chromatin loops can collide, a process that is negatively regulated by the cohesin release factor WAPL. Loop collision is further pronounced in WAPL knockout cells, suggestive of a “cohesin traffic jam”. In summary, single molecule multi-contact analysis methods can reveal how the myriad of regulatory sequences spatially coordinate their actions on individual chromosomes. Insight into these single allele higher-order topological features will facilitate interpreting the consequences of natural and induced genetic variation and help uncovering the mechanisms shaping our genome.

---

The invention of chromatin conformation capture (3C) technology^1^ and derived methods ^2^ has greatly advanced our knowledge of the principles and regulatory potential of three-dimensional (3D) genome folding *in vivo*. Insights obtained using these technologies include the discovery of regulatory chromatin loops that bring distal enhancers in close physical proximity to target gene promoters in order to increase their transcriptional output. These methods have also led to the identification of and architectural loops, often anchored by bound CTCF proteins, which form the structural chromosomal domains that spatially insulate transcription regulatory circuits^3,4^. Detailed topological studies and genetic evidence have further revealed that individual enhancers can contact and control the expression of multiple genes. Vice versa, single genes are oftentimes influenced by multiple enhancers^5,6^. Similarly, in population based assays individual CTCF sites can be seen contacting multiple other CTCF sites. Based on such observations it has been hypothesized that DNA may fold into spatial chromatin hubs ^7,8^. However, current population-based pair-wise contact matrices cannot distinguish clustered interactions from mutually-exclusive interactions that independently occur in different cells. To investigate the existence and nature of these hubs, high-throughput strategies are needed for robust detection, analysis and interpretation of multi-way DNA contacts.

Recently, several 3C procedures have been modified for the study of multi-way contacts, but most are limited in throughput and contact complexity ^9-12^ A recently developed genome-wide approach for multi-contact analysis, called C-Walks (chromosomal walks)^11^ gave an interesting glimpse of the spatial aggregation of genomic loci, indicating that cooperative hubs may be rare but present for example at polycomb bodies. C-walks, but also genome architecture mapping (GAM), another new method that analyzes genomic co-occurance frequencies in thin slices of fixed nuclei^13^, are difficult to scale up and currently don’t offer the resolution necessary to study higher-order topologies inside structural domains.

To enable the comprehensive study of spatial clustering of regulatory elements and genes, and dissect their interplay at the level of single alleles, we developed Multi-Contact 4C-seq (MC-4C). MC-4C is premised on the fact that 3C-based protocols generate aggregates of DNA segments that reside in each other’s 3D proximity in the nucleus. These ‘DNA hairballs’ are created via *in situ* formaldehyde-crosslinking of chromatin, followed by restriction enzyme-mediated DNA fragmentation and proximity-based re-ligation of crosslinked DNA fragments. The resultant DNA concatemers are characteristically sized >10kb^14^. Conventional 3C protocols trim these products further to enable efficient analysis of singular ligation junctions only. The MC-4C protocol is designed to keep these concatemers large, enabling the analysis of multi-way contacts for selected genomic sites of interest through third generation single molecule sequencing, such as Oxford Nanopore Technologies (ONT) minION. In brief, MC-4C entails the following steps. Like 4C-seq^15^ and Targeted Locus Amplification (TLA) technology^16^, MC-4C selectively PCR-amplifies concatemers with primers specific to a fragment of interest (the viewpoint). For this PCR to be sufficiently effective, 3C PCR template in the range of 2-5Kb is made by digesting the large concatemers with a 6-cutter restriction enzyme and re-ligation under conditions supporting self-circularization. To reduce prevalent rolling circle amplification and eliminate abundant non-informative undigested products, Cas9-mediated *in vitro* digestion of respectively the viewpoint fragment (in between the inverse PCR primers) and its two neighbor fragments is performed prior to PCR. After PCR, the product is size-selected (>1.5Kb) and sequenced on the MinION sequencing platform (**Fig. 1a**).

**Figure 1.**
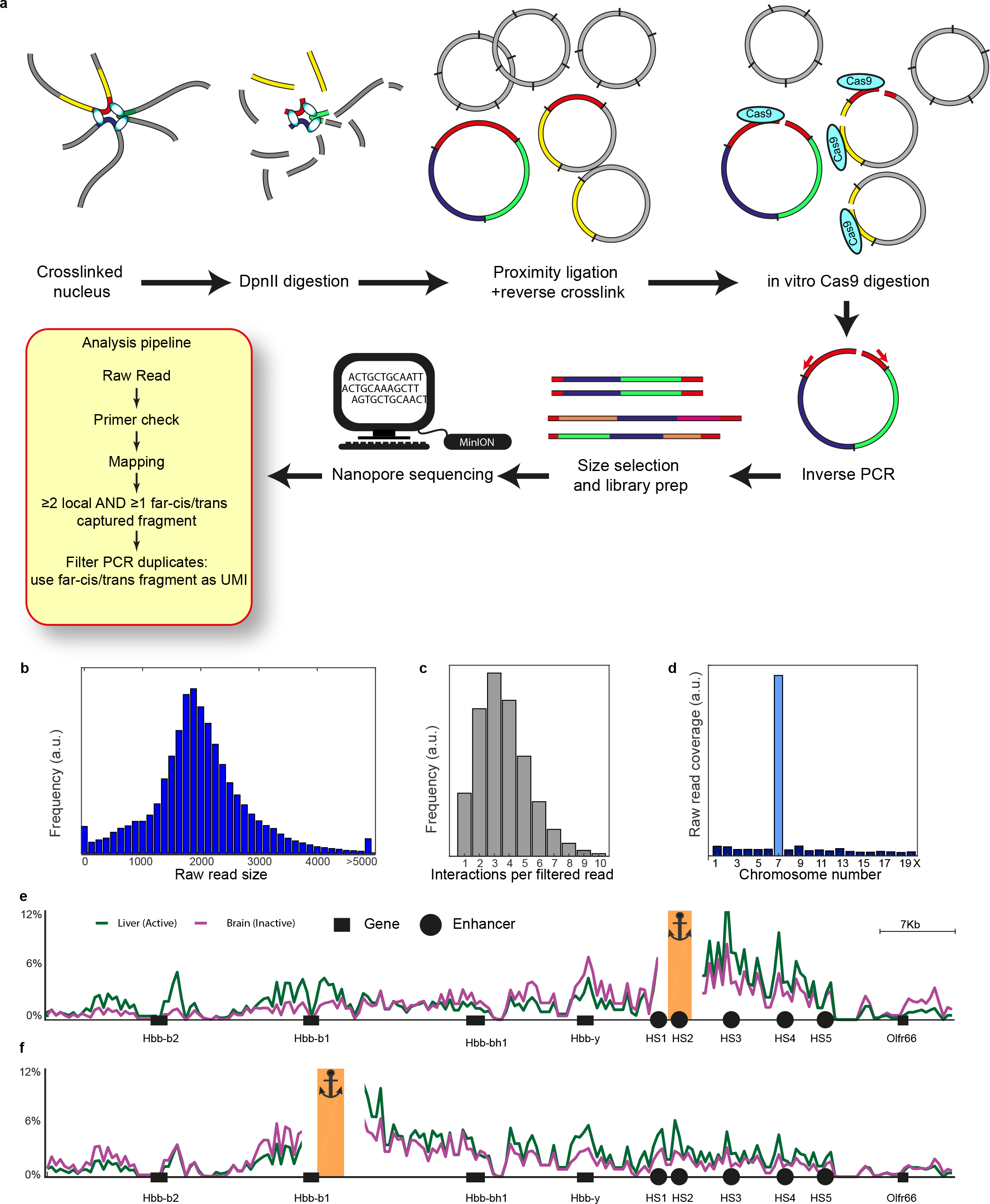
Multi-contact 4C technology. **a.** Schematic presentation of the MC-4C strategy. **b-d** statistics of the Hbb-b1 viewpoint in fetal liver cells. **b.** MC-4C read size distribution after ONT minION sequencing. **c.** The number of MC-4C captured fragments per allele. **d.** Chromosomal distribution of captured fragments. **e-f.** Overall (pan-allelic) MC-4C contact profile of β-globin HS2 (**e**) and Hbb-b1 (**f**) in E14.5 fetal liver (green) and brain (purple).

An integral component of MC-4C is its elaborate computational analysis strategy, which provides the necessary pre-processing of the ONT sequence data and downstream analysis to enable meaningful interpretation of allelic co-occurrence frequencies (See Suppl. Methods). To appreciate local multi-way contacts at the level of individual alleles, it is key to filter and select for the informative reads that have two or more contacts within a pre-defined chromosomal region of interest (ROI). In order to compute reliable statistics, it is also essential to efficiently remove all reads originating from PCR duplicates. In our method, PCR duplicate removal is guided by co-captured fragments far outside the ROI (see methods and **Supplementary Fig. 1a**): the chance of independently capturing such given fragment more than once is extremely small, implying that these sequences can serve as genomically contributed unique molecule identifiers (UMIs) in MC-4C. Because contact profiles are PCR filtered and thus a direct reflection of single allele measurements, they in principle become quantitative, albeit limited still by technically inherent variation that may arise from differences in cross-link, digestion, ligation and mapping ability between fragments.

To explore new biology that may be exposed from MC-4C we applied the technique to 3 different genetic systems: We chose the mouse β-globin and Pcdhα loci, both constituting multiple gene promoters and (super-) enhancer elements that act in concert to control defined developmental and cellular expression patterns. We also selected cohesin-looped topological domain boundaries that, upon cohesin stabilization, show extended loops with much more distal anchor sites in population-based Hi-C ^17^. We performed a total of 20 MC-4C experiments (27 minION sequencing runs) to obtain an average of 13K individual allelic micro-topologies, spanning an average total of 80K spatial contacts, per viewpoint (**Supplementary Table 1**).

**Figure 1** summarizes results from a typical MC-4C experiment. Because of PCR, which has a strong bias for small amplicons, and size selection, which we perform to remove the small amplicons prior to sequencing (see methods), the average size of MC-4C reads is approximately 2 kb (**Fig. 1b**, **Supplementary Fig. 2**). Most reads span 3-4 spatial contacts, some up to 10 (**Fig. 1c**, **Supplementary Fig. 3**), with spatial contacts being scored based on ligation events between restriction fragments that are not immediately juxtaposed in the reference genome. As in all other 3C methods, the great majority of captured sequences localizes to the immediate chromosomal vicinity of the viewpoint (**Fig. 1d**, **Supplementary Fig. 4**). Also, the contact profiles derived from sequences directly ligated to the viewpoint (i.e. those that one would analyze in ‘regular’ 4C-seq) are almost indistinguishable from those created from the indirectly ligated partners (**Supplementary Fig. 5**). Collectively this shows that the additional fragments that we capture and analyze by MC-4C are still the result of 3D proximity-based ligation events and represent topologically meaningful genomic multi-way contacts made with the viewpoint fragment.

We first studied higher-order conformations of the genetically well-characterized mouse β-globin locus. It carries two embryonic globin genes (Hbb-y and Hbb-bh1) that compete with two downstream adult globin genes (Hbb-b1 and Hbb-b2) for contacts with and activation by the upstream β-globin super enhancer (SE) during development. This SE, also known as the Locus Control Region (LCR), is composed of five regulatory elements (hypersensitivity sites (HS)1-5) of which HS1-4 show enhancer activity ^18^. Genetic studies in mice further demonstrated that the two developmentally distinct sets of genes compete for activation between sets, but not among members of each set, and that the four enhancer elements of the SE can compensate to a high degree for each other’s activity^19,20^. We performed MC-4C experiments in both fetal liver, where the adult genes are highly expressed and in fetal brain where the β-globin locus is transcriptionally silent. As viewpoints, we included Hbb-b1, HS2, HS5 (and HS3 exclusively in liver). When all fragments captured by the HS2 experiment are aggregated across all individually analyzed alleles in a, so-called, overall MC-4C contact profile, we find pronounced and precise interactions with the other SE constituents as well as with the active (mostly adult) gene promoters, specifically in expressing (fetal liver) but not in non-expressing (fetal brain) primary mouse cells (**Fig. 1e**). A similarly detailed and tissue-specific topology is appreciable from the overall MC-4C contact profiles that we obtained when using HS5, Hbb-b1 or HS3 as the viewpoint (**Fig. 1f**, **Supplementary Fig. 6**). MC-4C therefore recapitulates - with high precision - the previously observed conformational features of the β-globin locus^8,21,22^.

We next sought to analyze specific multi-way chromatin conformations adopted by the mouse β-globin locus. For this, we selected from each MC-4C dataset the allelic conformations that contain its viewpoint (VP) in contact with a given second site of interest (SOI), to then quantify and visualize the higher-order contact frequencies with the remaining co-occurring sequences. **Figure 2a** and **b** show two examples of such VP-SOI plots (see also **Supplementary Fig. 7**). As evidenced from the highly localized peaks exactly at the elements of the SE, alleles of the active β-globin locus that fold to have Hbb-b1 (**Fig. 2a**) or HS5 (**Fig. 2b**) in contact with HS2 are likely to also have the other constituents of the SE interacting. This is evidence for the existence of a regulatory enhancer hub. However, the mere detection of the other enhancer elements can just be the consequence of their random collision with the interrogated conformation.

**Figure 2.**
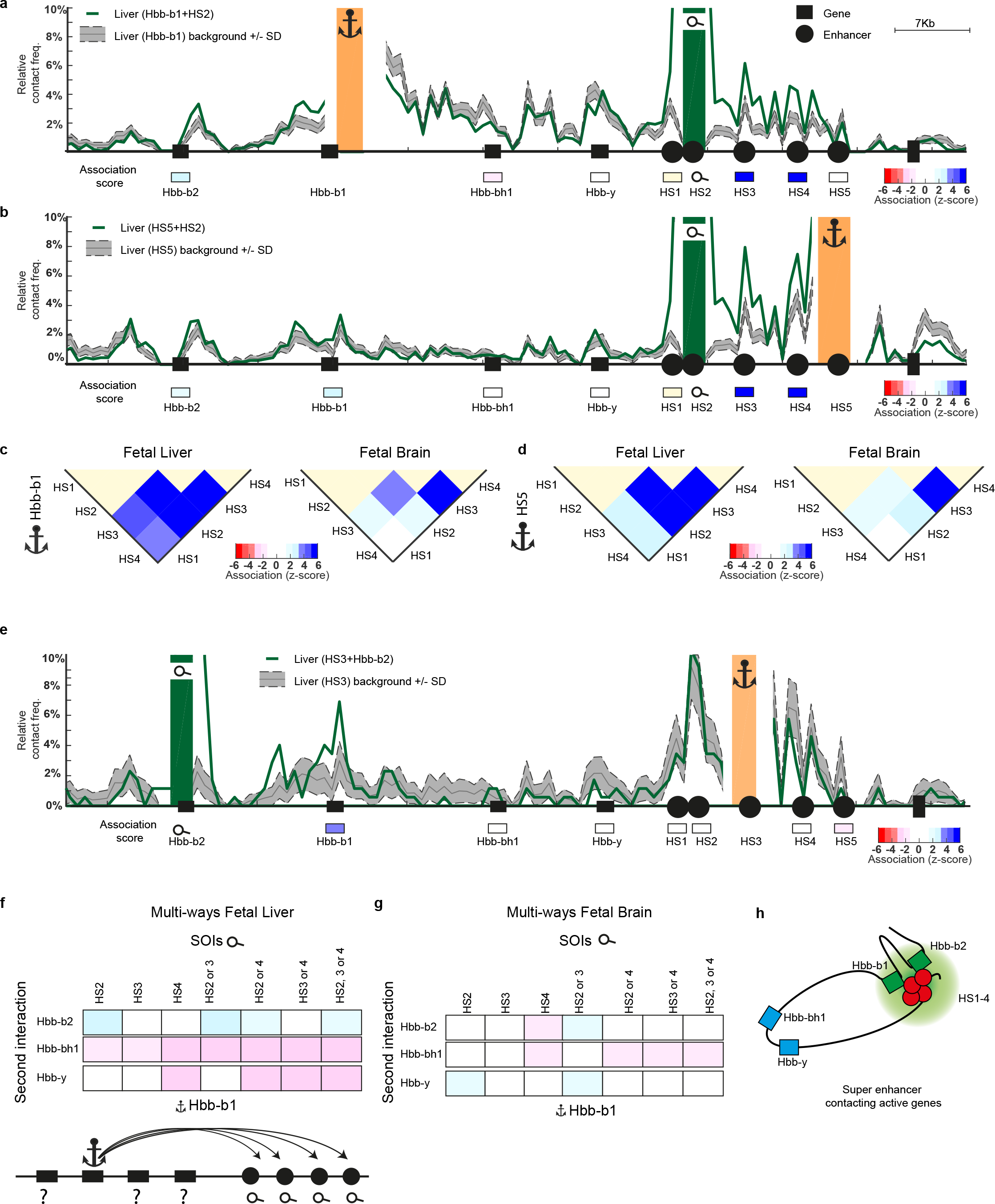
A β-globin super enhancer hub that can simultaneously accommodate two genes. **a-b.** Selected micro-topologies from fetal liver cells, having Hbb-b1 (**a**) or HS5 (**b**) in contact with HS2, are specifically enriched for the remainder constituents (HS3-4) of the β-globin super enhancer. Green line shows the observed and grey line the expected (−/+ SD) co-occurrence frequency of sites across the locus. Z-scores (dark blue indicating significant enrichment, dark red indicating significant depletion of a given site at the interrogated micro-topology) are shown for sites of interest in rectangles below each graph **c-d.** Summary of all z-scores for all possible pairs of SE elements (HS1-2 are too close to analyze interaction between), when one of them is in contact with the Hbb-b1 gene (**c**) or HS5 (**d**), in fetal liver (left: globin locus active) and fetal brain (right: globin locus inactive). Notice the preference for co-occurrence between non-neighboring SE elements specifically in fetal liver cells, revealing an active SE hub. **e.** Selected microtopologies from fetal liver cells, having HS3 in contact with Hbb-b2. Note that the other active gene, Hbb-b1, is preferentially found at these conformations. **f-g.** Summary of the clustering behavior (z-scores) of the three remainder globin genes when Hbb-b1 is in contact with each of the individual, or combinations of, SE constituents. **h.** Graphic showing the active β-globin super enhancer hub simultaneously contacting the two adult globin genes and excluding the two embryonic globin genes.

To distinguish cooperative from random or competitive multi-way interactions we plot the three-way co-occurrence frequency observed for a given VP-SOI conformation in terms of a z-score with respect to the distribution of contacts seen across all other conformations involving this VP (see methods and **Fig.2a** and **b** and **Supplementary Fig. 8**). Based on this analysis, we find that the individual elements of the β-globin SE are significantly enriched in conformations that already involve one of them. This preferred cooccurrence is appreciable in allelic conformations involving the distal downstream Hbb-b1 gene, as well as in those involving the upstream HS5 (**Fig. 2c**). To rule out that preferred co-occurrence is just a reflection of their linear proximity, we repeated the same MC-4C experiments on the same locus in non-expressing tissue (fetal brain). Here, no preferred multi-way interactions are observed, at least not beyond the directly neighboring constituents (**Fig. 2d**), showing that the spatial aggregation of β-globin SE constituents seen in expressing cells coincides with enhancer activity, not just linear proximity. We conclude that the individual elements of the active β-globin super enhancer can form a higher-order enhancer hub.

This super enhancer hub will be visited by the globin genes for their activation. To investigate the number of genes the hub can simultaneously accommodate, we analyzed the likelihood of Hbb-b2 and the two embryonic globin genes to be in contact with the SE when it is interacting with the adult Hbb-b1 gene (**Fig. 2f** and **g**). Despite their linear position in between the SE and Hbb-b1, the embryonic genes are clearly hindered in contacts with the SE when it is engaged with Hbb-b1, particularly in active tissue (**Fig. 2f** and **g**). This suggests that they physically compete with Hbb-b1 for interactions with the active enhancer hub. For Hbb-b2, the other adult globin gene that is more distal from the SE, we find no indication for physical competition with Hbb-b1 (**Fig 2e**). Its presence is either normally tolerated or even slightly stimulated in topologies having both SE elements and Hbb-b1 (**Fig. 2f**). MC-4C therefore provides original evidence for two novel higher-order topological phenomena. The first is that the individual elements of a single super enhancer, the active β-globin LCR, can cooperatively interact to form a spatial enhancer hub. The second is that this single enhancer hub can physically accommodate two genes at a time (**Fig. 2h**). We find that, in concordance with detailed genetic gene competition studies at this locus ^18-20^, partnering at the enhancer hub is allowed between developmentally synchronized genes, but not between genes active at different stages of development. The uncovered higher-order conformational features therefore provide a topological framework that helps interpreting the genetic observations.

In the mouse Protocadherin-Alpha (Pcdhα) gene cluster higher-order topologies may play a role in controlling its allelic expression patterns. Per allele, one of 12 alternative promoters (Pcdhα 1-α 12) is selected for expression. This ensures that individual neuronal cells express a unique repertoire of membrane-exposed protocadherin molecules, which is essential for axon avoidance^23,24^. Aside from the variable promoters, two constant promoters are present that are active in every neuron (Pcdhα C1 and Pcdhα C2). The activity of nearly all promoters is regulated by two downstream enhancers, HS7 and HS5-1 (only a C2 seems not to be influenced by HS5-1)^25,26^. Forward oriented CTCF binding to all promoters and reverse oriented CTCF binding to HS5-1 positively contributes to gene expression^27^. Alternative promoter methylation, which prevents CTCF binding, has been proposed to influence allelic promoter choice^28^. We designed viewpoint primers in both enhancers HS5-1 and HS7 and on the promoters of a 4 and a 11 (data from a 4 and a 11 and from HS5-1 and HS-7 were pooled due to the high similarity between overall profiles (**Supplementary Fig. 9**)) and performed MC-4C analysis in mouse E14.5 neuronal fetal brain cells, which express both Pcdhα variants (**Supplementary Fig. 9**), and in E14.5 fetal liver cells that do not express any of the Pcdhα promoters. All overall contact profiles showed that contacts between the enhancer and each of the promoter regions were perhaps slightly elevated in brain cells, but overall without dramatic differences in locus topology between fetal brain and liver cells. This suggests that there is no dominant, tissue-specific structure conserved in either fetal brain or liver cells (**Fig. 3a** and **b**). By selectively analyzing the allelic topologies having any of the enhancers in contact with a given alternative promoter in brain cells, we reasoned we could get insight into the specific folding of alleles expressing this particular alternative promoter. As an example, **Fig. 3c** shows how the other sequences of the locus participate in the micro-topologies centered around contacts between the α4 or a11 promoter, when these are in contact with HS7. In neurons, these configurations are specifically enriched for the other enhancer HS5-1 (39 kb downstream of HS7), as well as for the constitutively active Pcdh-α C2 promoter (73 kb upstream of HS7). In liver cells, the corresponding selection of micro-topologies do not specifically engage the HS5-1 enhancer, nor any of the genes, as expected if assuming that these contacts in non-expressing cells are a reflection of non-functional, random, collisions. The brain-specific enhancer hub involving cooperative interactions between HS7 and HS5-1 is similarly appreciable when studying other relevant subsets of allelic conformations (**Fig. 3d**). Additionally, α C2 is preferentially found at micro-topologies involving interactions between the enhancers and an alternatively transcribed Pcdhα promoter, while αC1 is not necessarily evicted from them. The Pcdhα active chromatin hub therefore appears capable of physically accommodating two or more genes at a time. We then wished to test whether physical competition for enhancer contacts between the α1-12 promoters may underlie their mutually exclusive allelic expression in neuronal cells. However, we found the α1-12 promoters being too close together on the linear chromosome template to observe such mutually exclusive contacts, at least at the current resolution of MC-4C (**Supplementary Fig.10**). In summary, as seen for the β-globin super-enhancer, the active linearly dispersed individual enhancers HS7 and HS5-1 and the Pcdh-αC2 gene promoter of the Pcdhα locus cooperatively interact to form a tissue-specific active chromatin hub that can simultaneously be contacted by at least one additional gene promoter (αC1 or α1-12). Importantly, our studies on Pcdhα further show that MC-4C can be used to characterize the interaction profiles of rare subpopulations of alleles, revealing topological features that are missed by population-based pair-wise contact analysis methods.

**Figure 3.**
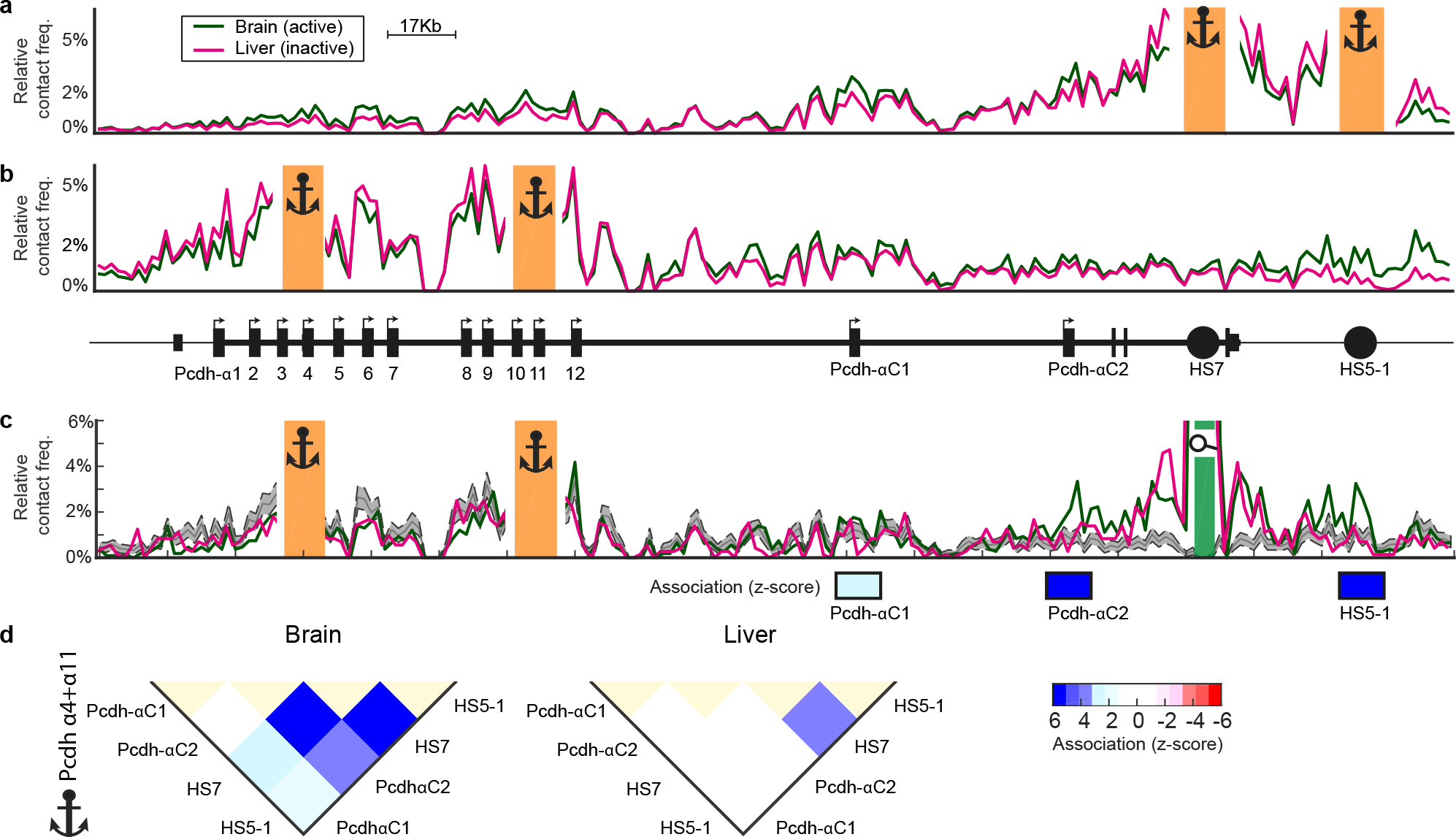
MC-4C uncovers Pcdhα hub conformations in tissue-specific sub-sets of cells. **a-b.** Overall (pan-allelic) MC-4C contact profiles of the combined HS7 and HS5-1 viewpoints (**a**) and the combined Pcdh-α4 and α11 viewpoints (**b**) in fetal brain (green: active) and fetal liver (purple: inactive). **c.** Selected micro-topologies from fetal brain (green: active) and fetal liver (purple: inactive), having Pcdh-α4 or α11 (the viewpoints, see anchors) in contact with HS7 (the SOI, see magnifying glass). In fetal brain these presumably are the rare allelic conformations that transcribe either Pcdh-α4 or α11. HS5-1 as well as the pan-cellularly active αC2 gene promoter preferentially cluster at these conformations. **d.** Summary of the clustering behavior (Z-scores) of the neuron-specifically active Pcdhα elements (αC1, αC2, HS5-1 and HS7) in fetal brain (left, locus active) and fetal liver (right, locus inactive) when any one of these elements is in contact with either Pcdh-α4 or αll. Notice the preference for co-occurrence between αC2, HS5-1 and HS7, specifically in the active Pcdhα locus (brain).

As a third model system to study multi-way chromatin interactions we focused on CTCF/cohesin-anchored chromatin loops. Cohesin is a ring-shaped protein complex that is necessary to form loops between CTCF-bound domain boundaries^29,30^. The ‘loop extrusion’ model^31,32^ predicts that cohesin forms loops by an active process in which the chromatin fiber is pulled through its lumen. The loop is then processively enlarged until two compatible roadblocks, e.g. two convergently oriented and occupied CTCF sites, are reached, at which the loop is stably anchored. Without WAPL, cohesin remains bound to chromatin for longer periods of time, which enables given CTCF sites to engage with new CTCF partners over much larger distances, as measured by population Hi-C across the population of WAPL deficient Hap1 cells (ΔWAPL) cells^17^. One possibility is that these additional ultra-long range interactions are the result of cohesin progressing beyond original CTCF roadblocks to mediate direct pairing between more distal CTCF sites. An alternative explanation would be that distant sites are reeled in through the aggregation of CTCF loop anchors (loop ‘collision’), which ultimately brings together distal CTCF sites. Population-based pairwise contact studies cannot distinguish between these two scenarios. MC-4C, which allows quantification of allelic co-occurrence frequencies, does enable unentangling of these two scenarios.

We selected a region that clearly showed novel long-range contacts in ΔWAPL cells based on Hi-C data (**Fig. 4a**) and applied MC-4C to two CTCF sites that anchor these loops. A comparison between their pan-allelic contact profiles in WT and ΔWAPL cells confirms that MC-4C also identifies these long-range contacts specifically in the ΔWAPL cell population (**Fig. 4b**). If they occur due to the skipping of CTCF roadblocks, we would expect a severe depletion of intervening CTCF sites from the allelic micro-topologies having these distal CTCF sites together. We find the opposite: intervening CTCF sites show a strong preference to aggregate with these structures, something we observe irrespective of the combination of new long-range contacts we interrogate at this locus (**Fig. 4c-d** and **Supplementary figure 11**). To exclude that we are identifying locus-specific effects, we applied MC-4C to another locus showing profound new contacts between distal CTCF sites in ΔWAPL cells. Also here we find no evidence for mutual exclusivity between CTCF sites that at the cell population level all seem to interact with each other. Instead, as seen before, they are preferentially found clustered at single alleles (Supplementary Fig. 11 **and 12**). Therefore, rather than -or at least in addition to- the skipping of CTCF roadblocks, our data strongly suggests that WAPL depletion results in ‘loop collision’, with distal CTCF sites getting into contact because of progressive aggregation of loop domain anchors. Based on Hi-C it was also noted that, in the absence of WAPL, contacts between ‘illegal’ (non-convergent) oriented CTCF sites are more frequently observed^17^. This now seems partially explained as an inevitable result of cluster formation: when three or more CTCF sites form topological aggregates, at least one must be in the ‘ wrong’ orientation.

**Figure 4.**
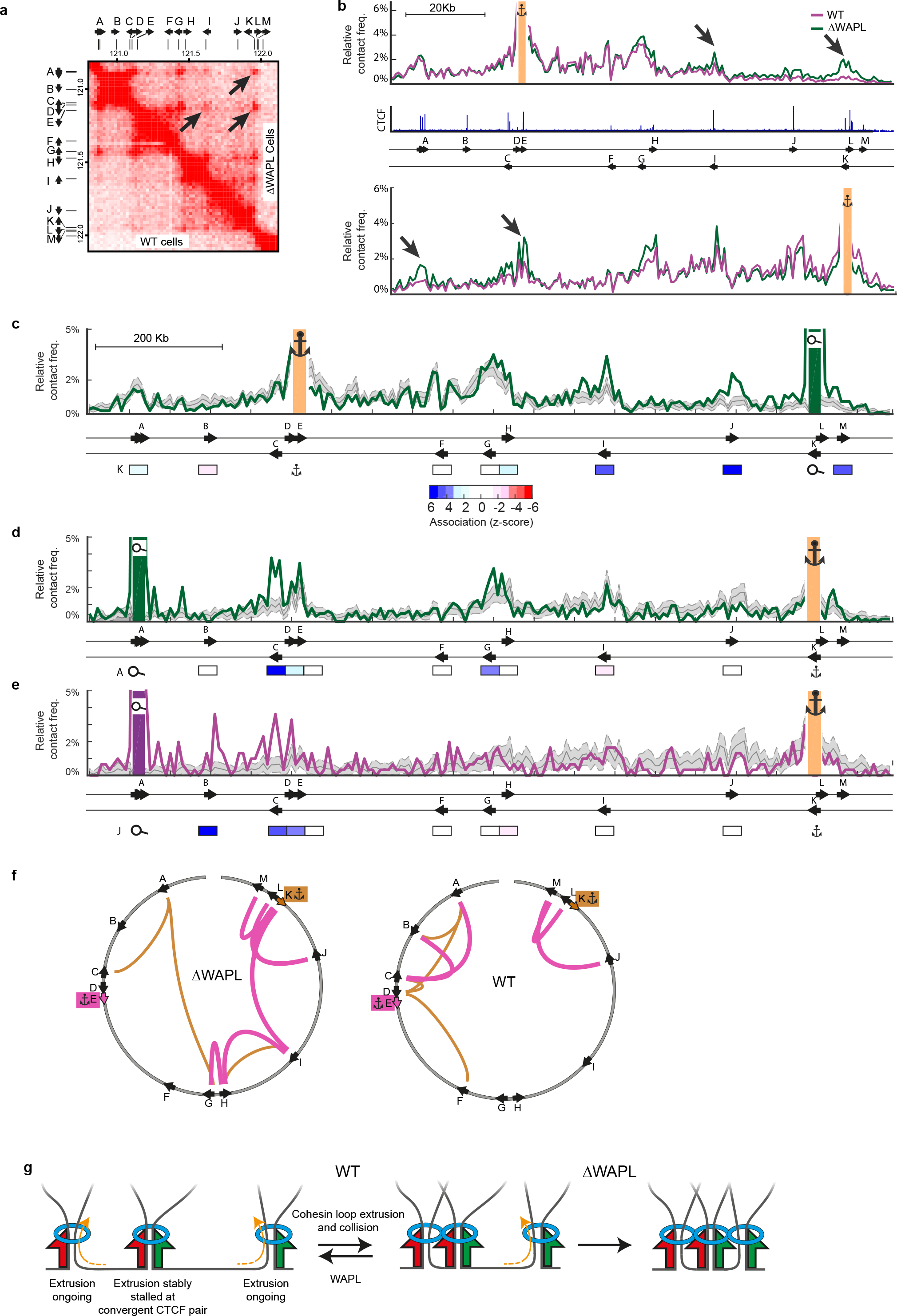
Depletion of WAPL stimulates collision of CTCF-anchored domain loops. **a.** Hi-C contact matrix of a selected genomic region in wild-type (upper right part) and ΔWAPL (lower left part) HAP-1 cells. Position and orientation of CTCF-binding sites are indicated. Arrows point at new, long-range, contacts that appear upon WAPL knockout. **b.** Overall (pan-allelic) MC-4C contact profiles of an upstream, forward oriented, CTCF site E (top) and a downstream, reverse oriented CTCF site K (bottom), in WT (purple) and ΔWAPL (green) HAP1 cells. CTCF ChlP-seq profile (from WT) is shown in blue, with the orientation of CTCF sites indicated below the ChlP-seq track. **c.** Selected micro-topologies from ΔWAPL cells, having CTCF site E (Fw) in contact with CTCF site K (Rv). The z-scores are plotted below, showing preferred clustering of CTCF site I and J at this conformation. **d.** Selected micro-topologies from ΔWAPL cells, having CTCF site K (Rv) in contact with CTCF site A (Fw). The z-scores are plotted below, showing preferred clustering of CTCF site C and G at this conformation. **e.** Selected micro topologies in WT HAP1 cells, having CTCF site K (Rv) in contact with CTCF site A (Fw). With z-scores plotted below, indicating that the rare allelelic conformation where K interacts with with A, co-occurs with interactions with B, C and D, but not with any of the CTCF sites in-between K and D. **f.** Preferential contacts propensity between CTCF sites. Links are colored with respect to view point and their thickness depict strength of preferential contacts between CTCF sites for KO vs. WT. **g.** Proposed “traffic jam” model explaining the increased incidence of CTCF cluster formation in ΔWAPL cells.

Since WAPL serves to destabilize, but not to prevent, loop formation, we reasoned that loop anchor clusters may also exist, albeit less frequently, in WT cells. To investigate this, we selected alleles from WT cells that had the same long-range CTCF contacts interrogated earlier in ΔWAPL cells. Importantly, these interactions are too rare in WT cells to stand out in population-based Hi-C and pan-allelic MC-4C contact profiles (**Fig. 4a** and **b**). Strikingly however, also in WT cells these now rare allelic conformations show a strong enrichment of intervening CTCF-based loop anchors. When we quantify the percentage of alleles showing simultaneous clustering of three or more distinct CTCF anchors, we see an increase from 5,6% to 8,6% (for the downstream viewpoint) and from 6,8% to 10,9% (for the upstream viewpoint) in ΔWAPL as compared to wildtype cells. We therefore conclude that loop collision and anchor aggregation also occurs in wild-type cells but less frequently, due to the counteracting effect of WAPL (**Fig. 4e** and **f**, **Supplementary Fig. 11**). In light of the loop extrusion model, our findings could be explained by assuming a “cohesin traffic jam”. Any cohesin ring that is extruding a DNA loop (or sliding over the DNA strands) will at some point be released from DNA by WAPL. If not, it will inevitably encounter and presumably be stopped by another cohesin ring that was already immobilized at a CTCF roadblock. Subsequent cohesin rings could then start reeling in other CTCF sites from both directions or as nested loops (loops within larger loops), eventually leading to the spatial aggregation of CTCF-bound loop anchors. Collisions from inside and outside an existing loop would then result in a “cohesin traffic jam” (**Fig. 4g**). Although just a theory, loop collisions resulting in a “cohesin traffic jam” fits well with the high frequency of illegal loops seen in ΔWAPL cells, but also with the “vermicelli” cohesin staining patterns observed in ΔWAPL cells^17,33^.

In summary, we present MC-4C which allows for high resolution analysis of spatial DNA sequence co-occurrence frequencies at the single allele level. MC-4C contact counts represent relative, not absolute, contact frequencies, as one cannot assume that not being captured (i.e. not being crosslinked, digested, ligated and mapped to the genome) does imply not being together. We present a method that allows statistically distinguishing cooperative from random and competitive interactions, for chosen genomic regions. The data show that by this method sequences that directly neighbor each other on the linear chromosome are being scored as obligatory together in 3D space (cooperative interactions). This is not only as expected (physically connected sequences simply cannot spatially escape each other), it can also be biologically meaningful: it is not without reason that only when transcription factor binding motifs cluster on the linear chromosome they can form functional regulatory motifs. It does emphasize though that, for correct interpretation of MC-4C results, resolution must be high enough to discern spatial clustering as the mere consequence of linear physical proximity from that driven by biological processes. Here, we accomplish this by analyzing often more than ten thousand independent allelic conformations per experiment and by comparing allelic co-occurrence frequencies of the same locus in its active versus inactive configuration. The study of higher-order chromatin topologies at such high resolution uncovers new biology: individual elements of a super enhancer aggregate to form an enhancer hub that can simultaneously accommodate multiple genes at a time. Similarly, we also find that cohesin drives aggregation of CTCF-bound domain boundaries, which is counteracted by WAPL. The latter studies, as well as our work on Pcdhα, further demonstrate that MC-4C can identify and analyze relevant structures missed by population-based contact methods like Hi-C or 4C-seq because they are present only in a small percentage of cells. High resolution multi-way contact analysis methods like MC-4C promise to uncover how the multitude of regulatory sequences and genes truly coordinate their action in the spatial context of the genome.

For the visualization of co-occurrence frequencies of any site of interest with a given MC-4C viewpoint, and the calculation of the significance of such three-way interactions, we refer to the interactive viewer that we made available, together with the data shown in this manuscript at: <u>www.multicontactchromatin.nl</u>.

## Methods

### Tissue and cell culture

Mouse embryos were harvested at 14.5 days post conception, livers and brains were manually dissected. Cells were brought into single-cell suspension in 10% FBS/PBS using a 40μM strainer. Wild-type Hap1 cells and WAPL KO Hap1 cells were cultured and harvested as described in Haarhuis *et al.*

### MC-4C template preparation

MC-4C template was prepared following the regular 4C protocol (described in van de Werken *et al.* and Splinter *et al.*), with several adjustments. DpnII (liver and brain) or MboI (Hap1 cells) digestion was performed in a 500μL volume and the first ligation was performed in a 2 mL volume. After reverse crosslinking, DNA was precipitated using 20μL NucleoMag P-beads (Macherey-Nagel) and 2 mL 2-Propanol, washed twice using 80% ethanol and resuspended in 1x restriction buffer appropriate for the secondary digestion (HindIII in all cases except for the Man1A viewpoint, where SacI was used). After overnight digestion with the second restriction enzyme, the enzyme was heat inactivated and the template was circularized by diluted ligation (5ng/μL DNA). After ligation the DNA was purified using P-beads (l0 μL per mL of ligation volume), and two wash steps using 80% ethanol. To remove any remaining P-beads, the template was purified using the Quiagen PCR purification kit.

### *In vitro* Cas9 digestion of MC-4C template

Per viewpoint, three sgRNAs were designed using the ATUM online design tool: one on each flanking fragment and one in-between the viewpoint primers. gRNA *in vitro* transcription template was made using a PCR with two partially overlapping primers (as described in Nakayama *et al.^34^*). In vitro transcription was done using the megashortscript T7 transcription kit (Invitrogen). RNA was purified using 4x AMPure purification (Agencourt) while using DEPC water and avoiding RNAse contamination. Purified Cas9 protein was kindly provided by P. Shang and N. Geijsen. Cas9 was pre-incubated with the appropriate sgRNA. For typical experiments, we pre-incubated in a 300μL volume combining 600ng of gRNA with l5 pmol of Cas9 protein for 30 minutes at room temperature. Subsequently, all preincubations were added to 20μg MC-4C template DNA and incubated for 3-6 hours at 37°C for digestion. After the digestion, Cas9 was inactivated by adding l/25^th^ volume of l0% SDS and incubation at 70 °C for 5 minutes. The template was subsequently purified using a 0.6X AMPure purification.

### MC-4C viewpoint specific PCR

Inverse MC-4C primers were designed on DpnII-DpnII fragments, with an approximately 50bp offset from the restriction sites (where possible), to facilitate viewpoint detection in the analysis pipeline. PCRs were performed in 96 separate 25uL reactions, using l00ng of MC-4C template per reaction. PhireII polymerase (Thermo Fisher) was used with the following protocol: 98° for 30 seconds, followed by 3l cycles of: 98° for l0 seconds, 57° for 20 seconds and 72° for l minute and 30 seconds, followed by 72° for five minutes. After PCR amplification, all reactions were pooled and 1mL was simultaneously purified and size selected using a 0.6x AMPure beads.

### Pcdh-α expression analysis

RNA was isolated from fetal brain cells using TRIzol (Thermo Fisher), cDNA was produced using MMLV reverse transcriptase (Promega) and random primers (Promega). PCR was performed on cDNA using the primers described in Kaneko *et al.*^35^

### MinION library preparation and sequencing

Pippin HT size selection within a l.5-8kb range was performed on PCR products. Subsequently, libraries were prepared using the Oxford nanopore sequencing kits and sequenced with appropriate flow cells.

### MC-4C mapping and association analysis

Long Nanopore reads are split into fragments based on the restriction enzyme sequences in the reference genome and mapped. Due to the relatively high error rate in Nanopore reads, restriction sites may have been missed or erroneously introduced. To mitigate this, we employ a split-read capable aligner BWA-SW that is able to further split a fragment (in case of a missed restriction site). We additionally merge fragments mapped adjacent (in case of a spurious restriction site). In case fragment-ends do not coincide with restriction sites, fragments are shrunk in case their fragment-end is within l0 bp of a restriction site, otherwise they are extended to the next restriction site in the reference genome.

To remove PCR duplicates we utilize the fact that fragments that map far away from the VP are likely to be the result from random ligations and should therefore occur only once. Such fragments can thus be used as a UMI, and circles that share one or more UMI are filtered as duplicate, retaining only the one with the most fragments mapped to the locus of interest. Finally, circles with fewer than two fragments in the locus of interest are discarded.

To assess preferential contacts between the VP and two loci of interest (say X and Y), the set of circles that contain fragments in locus X (positive set) are compared to the set of circles that are devoid of fragments in locus X (negative set). To this end, we bootstrap the negative set to the size of the positive set and repeat this process l000 times to obtain the mean and standard deviation of the contact frequency between VP and Y in the absence of X. To prevent bias in the number of fragments per circle between the positive and negative set that is due to the fact circles in the positive set are required to contain a fragment in locus X, we randomly remove a fragment from the circles in the negative set. A z-score is calculated to express the significance of the observed associations.

## Supplementary Figure Legends

**Supplementary Figure 1.**
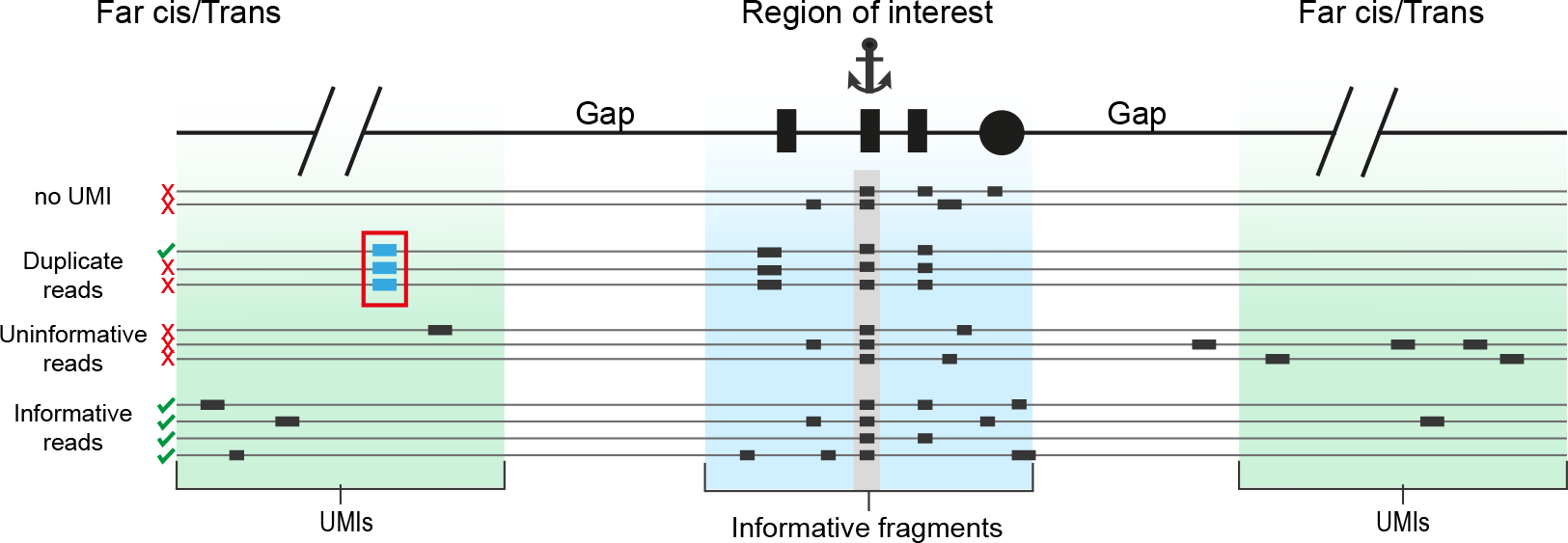
Schematic overview of critical data pre-processing steps. Scheme shows the removal of (1) reads with less than two captured fragments in the region of interest (ROI), (2) reads with less than one far-cis/trans fragment and (3) PCR duplicate reads, as guided by the capture of identical far-cis/trans fragments (genomically contributed unique molecule identifiers (UMIs)).

**Supplementary Figure 2.**
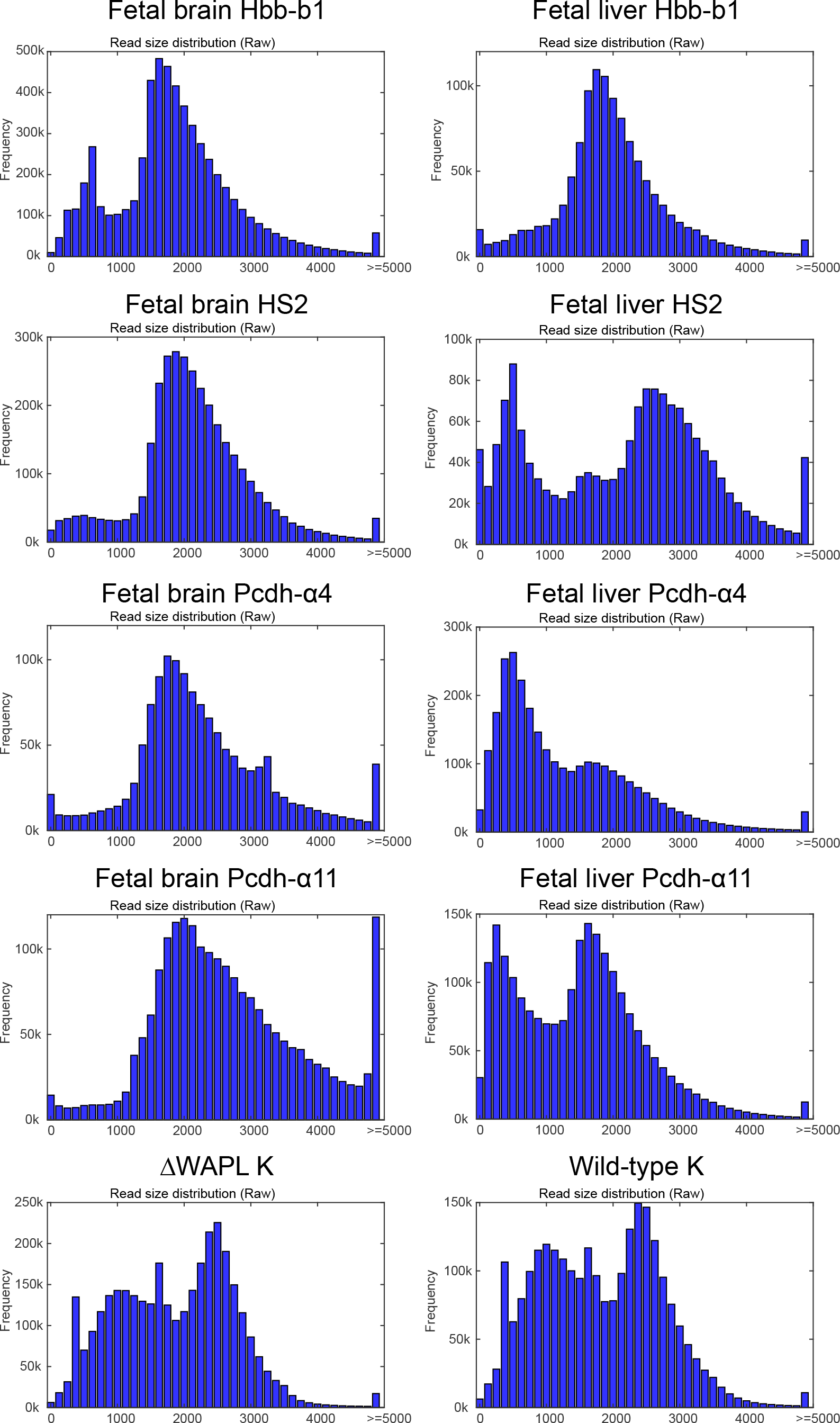
MC-4C read size distribution, after filtering. Read size distribution plots of ten representative MC-4C experiments, using viewpoints inside the α-globin and Pcdhα locus in primary mouse fetal liver and fetal brain cells and viewpoints at CTCF sites in wildtype and WAPL knockout human HAP1 cells.

**Supplementary Figure 3.**
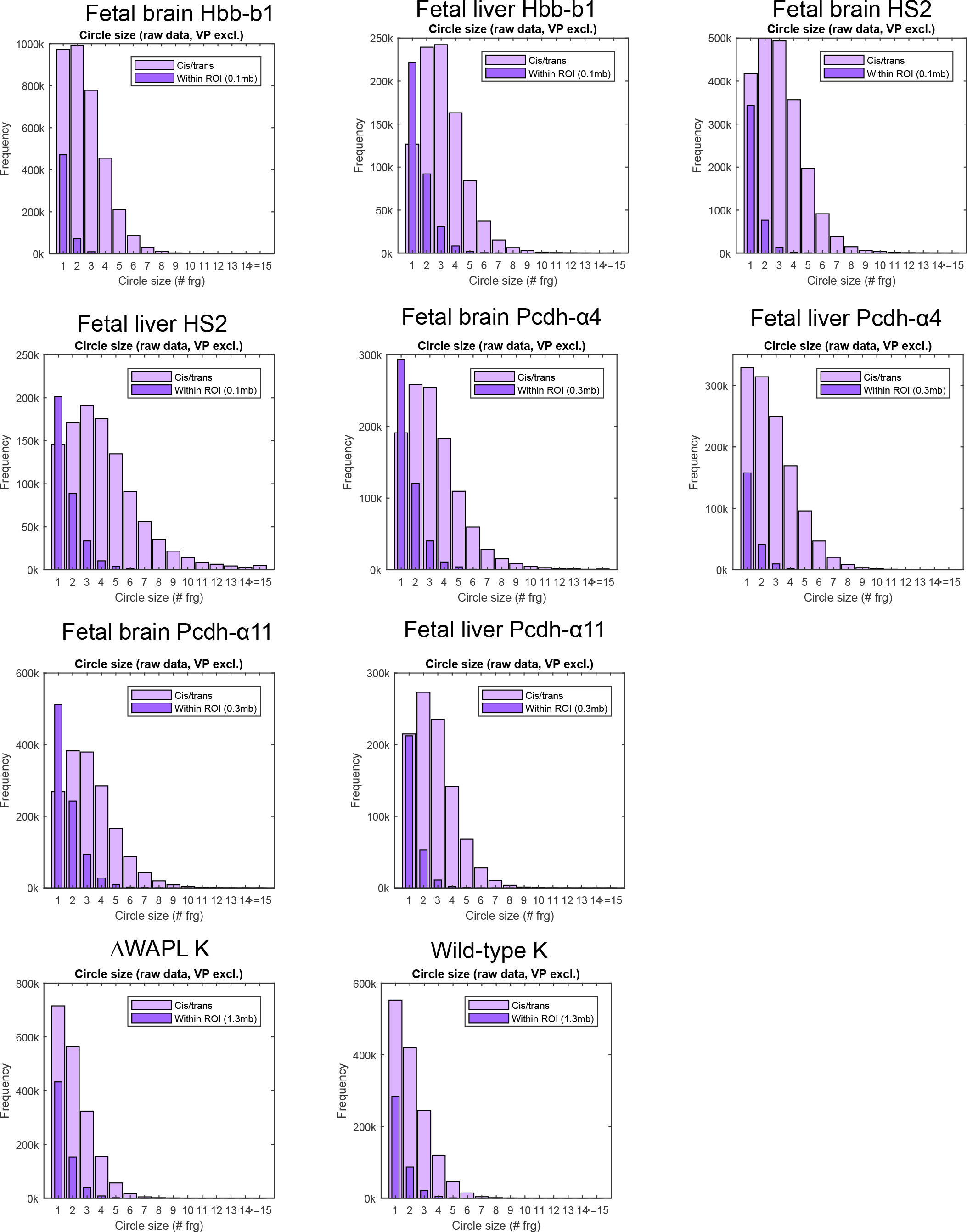
Number of captured fragments per read. Plots show number of captured fragments (i.e. number of identified contacts) per read, for representative MC-4C viewpoints inside the β-globin and Pcdhα locus in primary mouse fetal liver and fetal brain cells and viewpoints at CTCF sites in wildtype and WAPL knockout human HAP1 cells. Note that restriction fragments which map immediately next to each other on the reference genome are together counted as a single fragment (i.e. single contact).

**Supplementary Figure 4.**
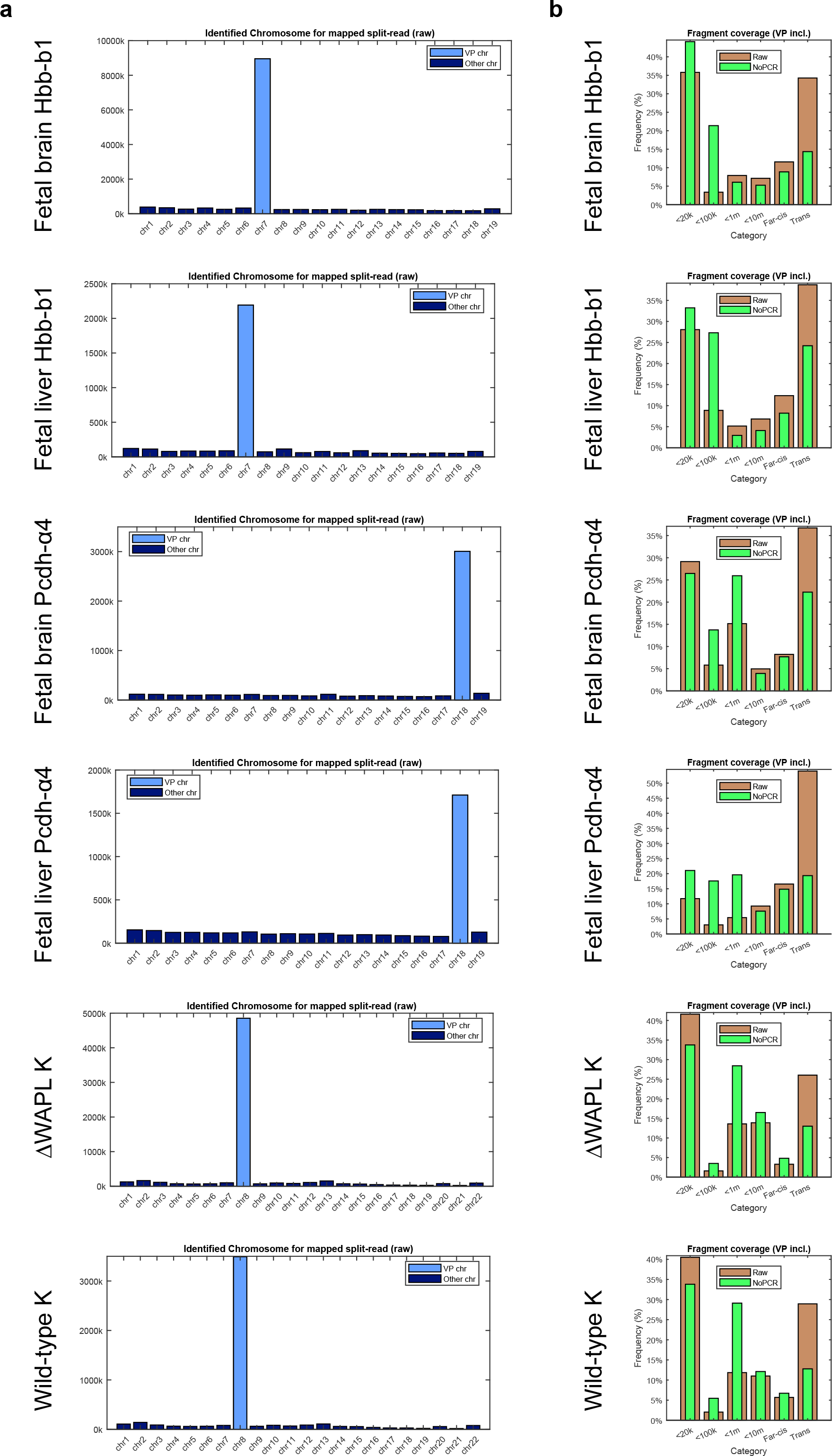
Genomic distribution of MC-4C captured fragments. **a.** Chromosomal distribution of captured fragments for ten representative MC-4C experiments. **b.** Distribution of captured fragments across chromosomal intervals at increased distance from the viewpoint, for ten representative MC-4C experiments.

**Supplementary Figure 5.**
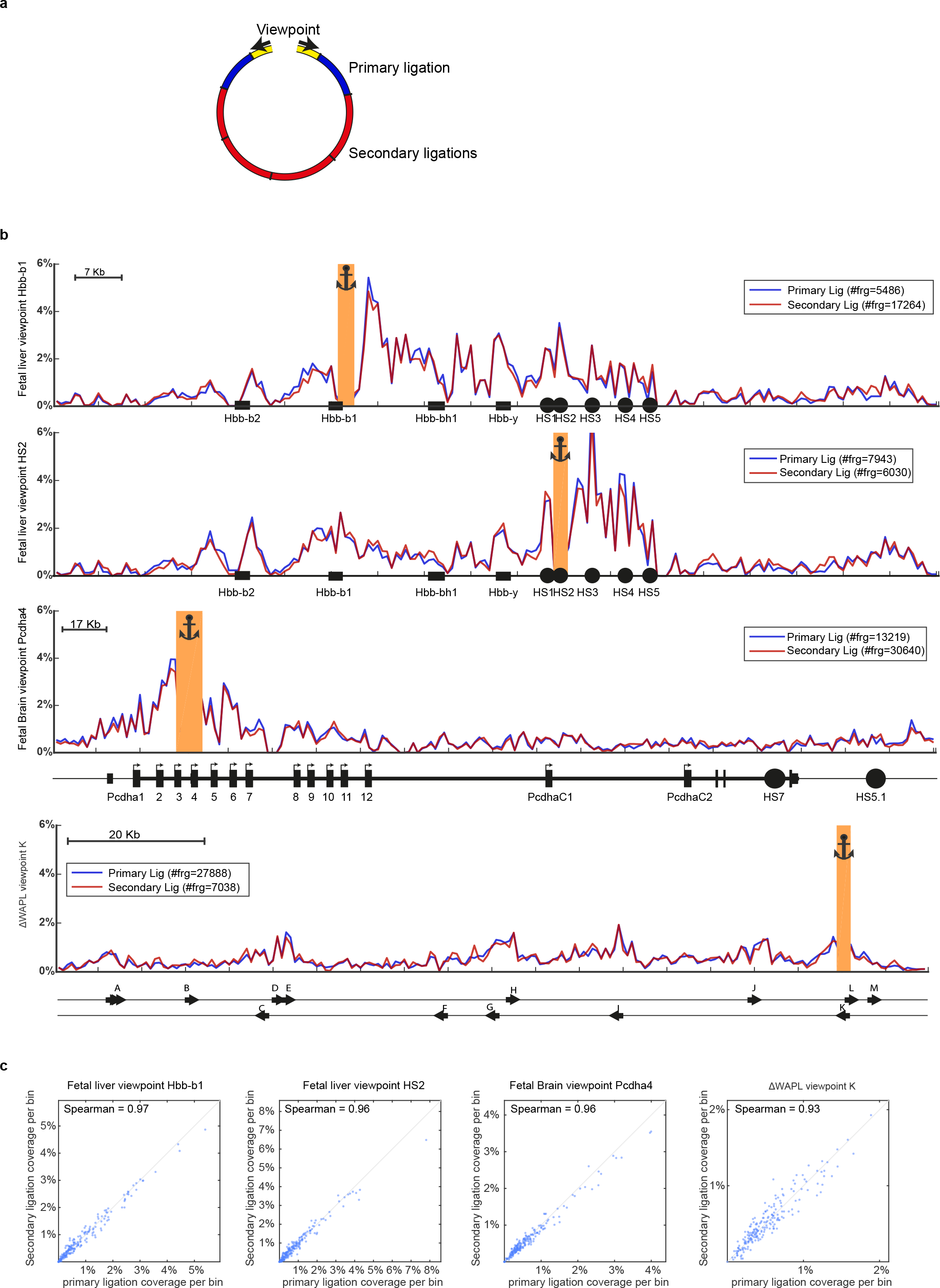
Similarity between primary and secondary ligation products. **a.** Cartoon explaining the difference between primary ligation products (as analyzed by Hi-C and 4Cseq) and the secondary ligation products that are additionally captured, sequenced and analyzed in mC-4C. **b.** Overlay of pan-allelic contact profiles of primary and secondary ligation products, for four representative MC-4C experiments. **C.** Comparison of the distribution of primary and secondary captured fragments across chromosomal intervals at increased distance from the viewpoint, for ten representative MC-4C experiments.

**Supplementary Figure 6.**
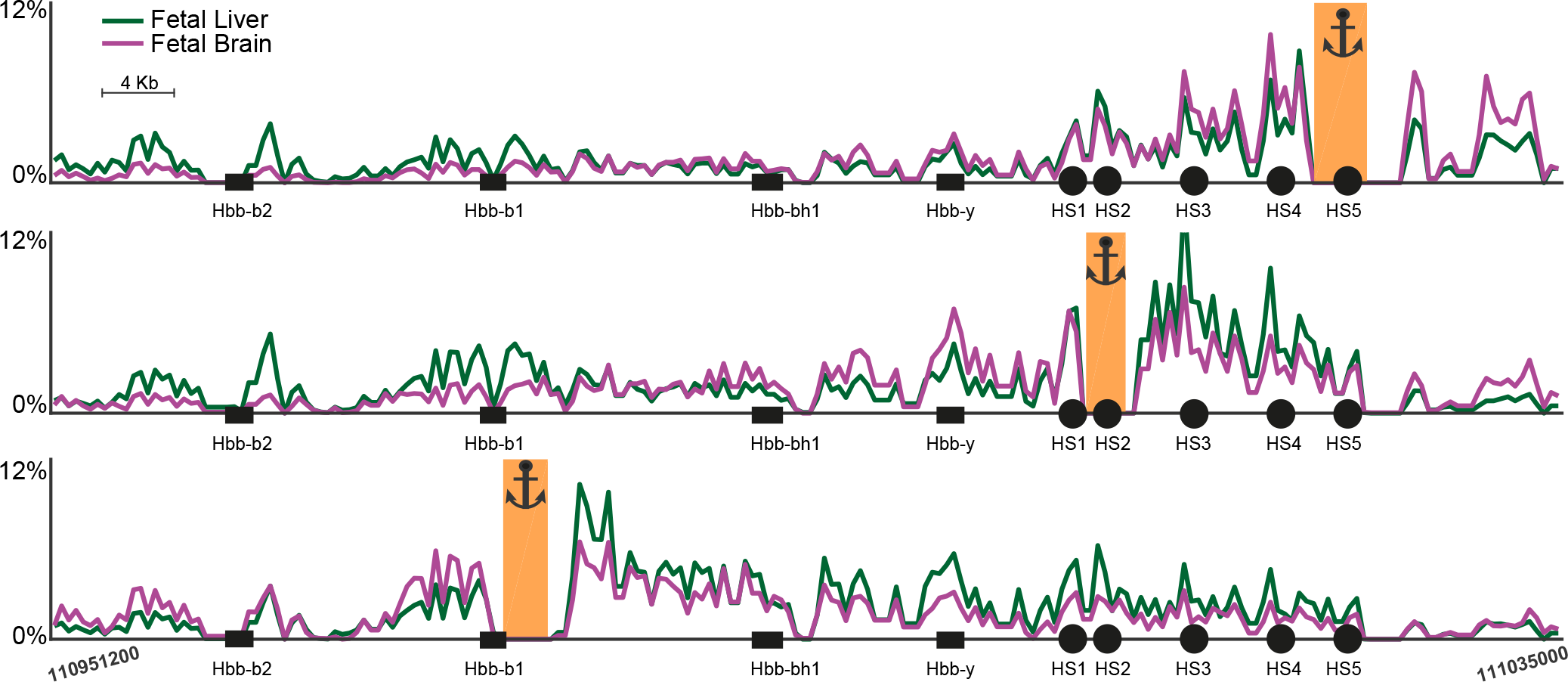
Overall (pan-allelic) MC-4C contact profiles of HS3 and HS5, in E14.5 fetal liver and fetal brain cells. In E14.5 fetal liver, the hbb-b1 and hbb-b2 genes are the predominantly active globin genes, while hbb-y expression is being silenced. In fetal brain cells, all globin genes are silenced (but residual expression may come from contaminating circulating blood cells).

**Supplementary Figure 7.**
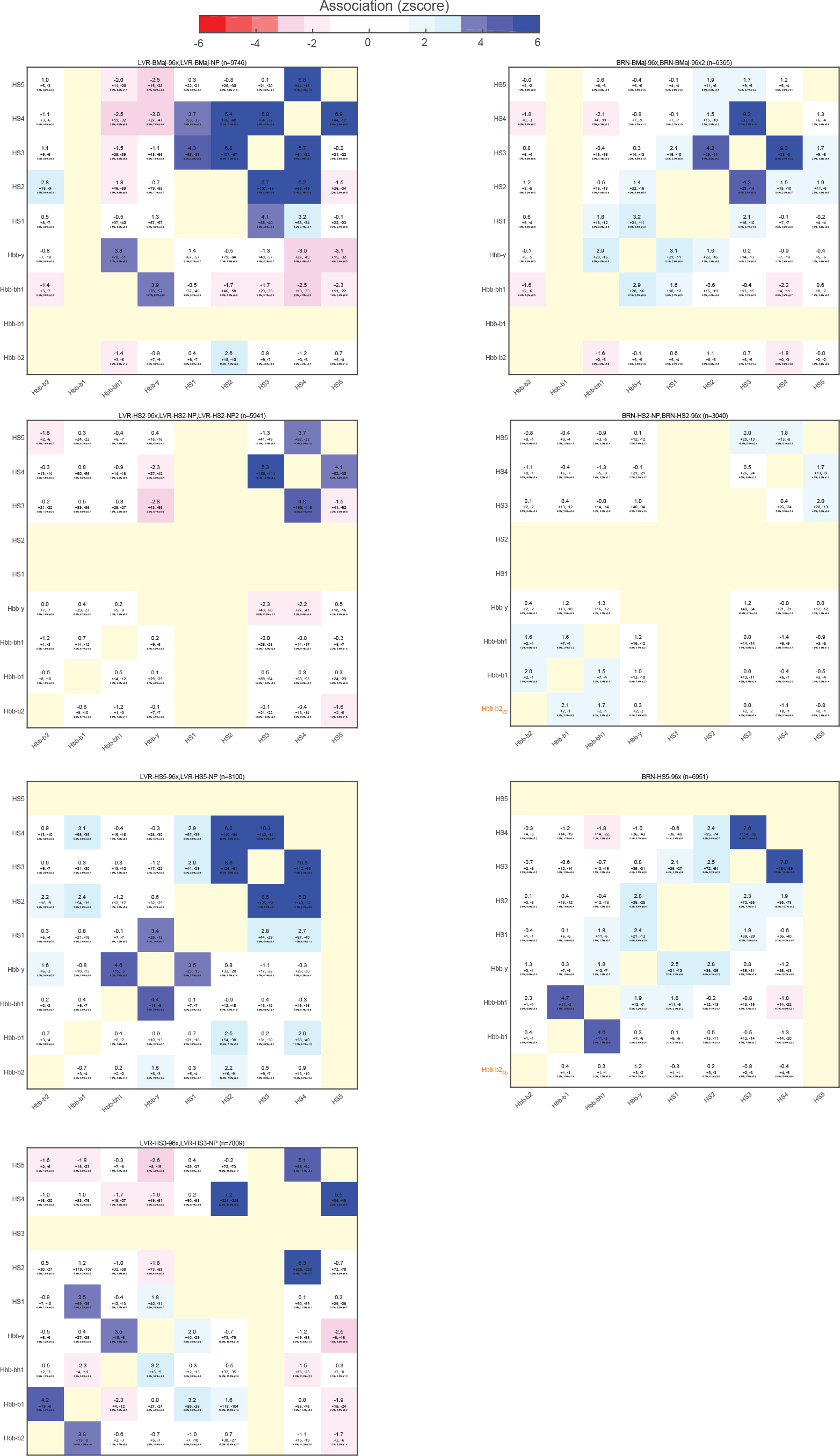
Allelic co-occur frequencies at the active and inactive β-globin locus. Complete overview of co-occur significance scores found between any pair of genes and/or super enhancer constituents in experiments using Hbb-b1, HS2, HS3 and HS5 as viewpoints, in E14.5 fetal liver and E14.5 fetal brain cells.

**Supplementary Figure 8.**
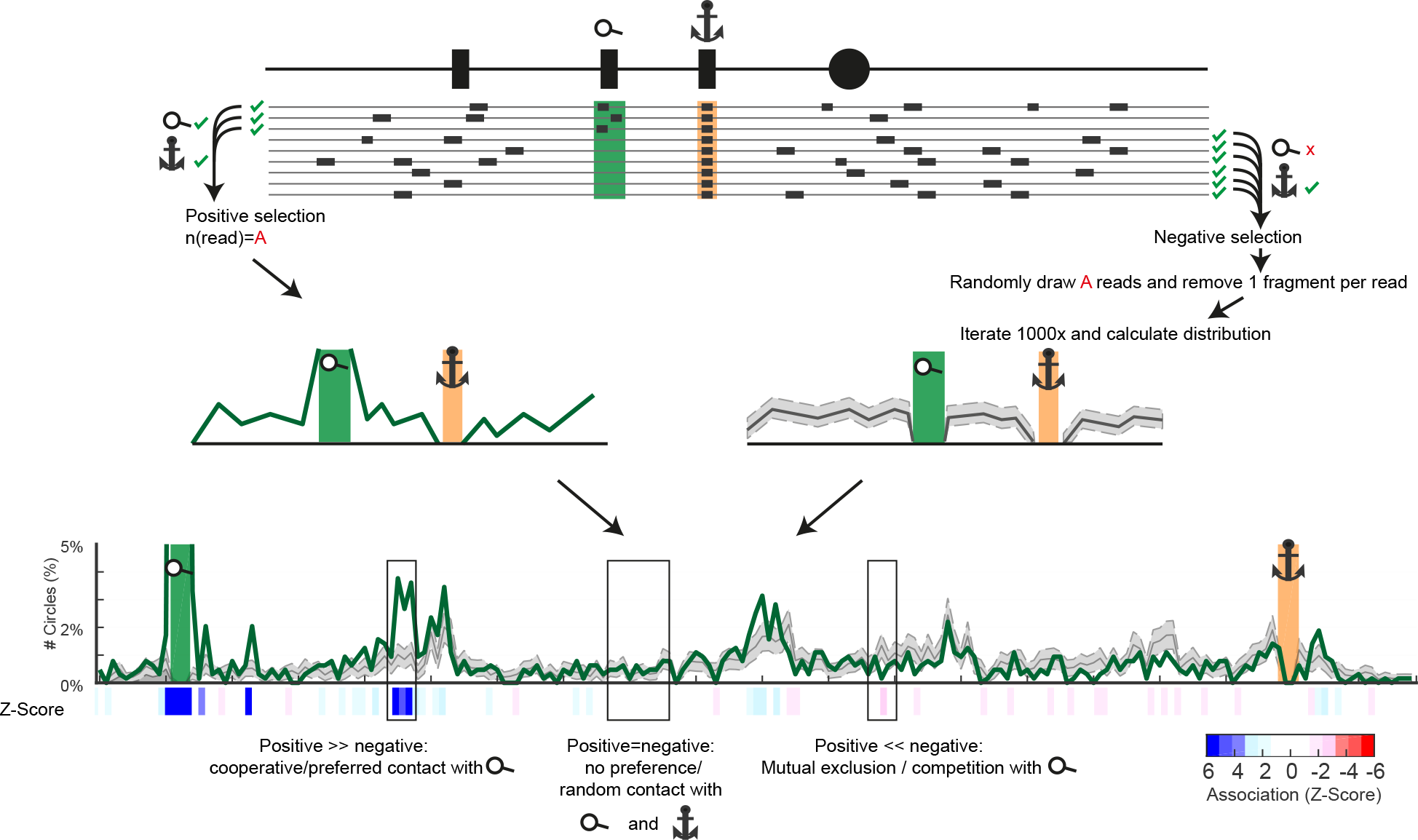
Distinguishing cooperative from random and competitive DNA interactions. To understand the relevance of any third sequence being co-captured at its observed frequency at micro-topologies involving a given VP-SOI contact, this frequency is compared to that expected from all other topologies centered on the VP but NOT involving this SOI. For this, all positive (VP-SOI containing) allelic topologies are separated from the large pool of remainder VP-centered allelic topologies (negative selection). Iterative (l000x) random drawing of an X number of alleles from this negative selection (with X being the exact number of VP-SOI containing alleles) is then performed to estimate the expected frequency distribution of a third partner. Based on z-scores we can then calculate whether this given third partner is significantly enriched in the positive set of alleles (which implies cooperativity with the SOI), significantly depleted from the positive selection (indicative of the third partner competing with the SOI for contacts with the VP) or equally distributed between the positive and negative selection (showing its random co-occurrence at the interrogated VP-SOI topologies).

**Supplementary Figure 9.**
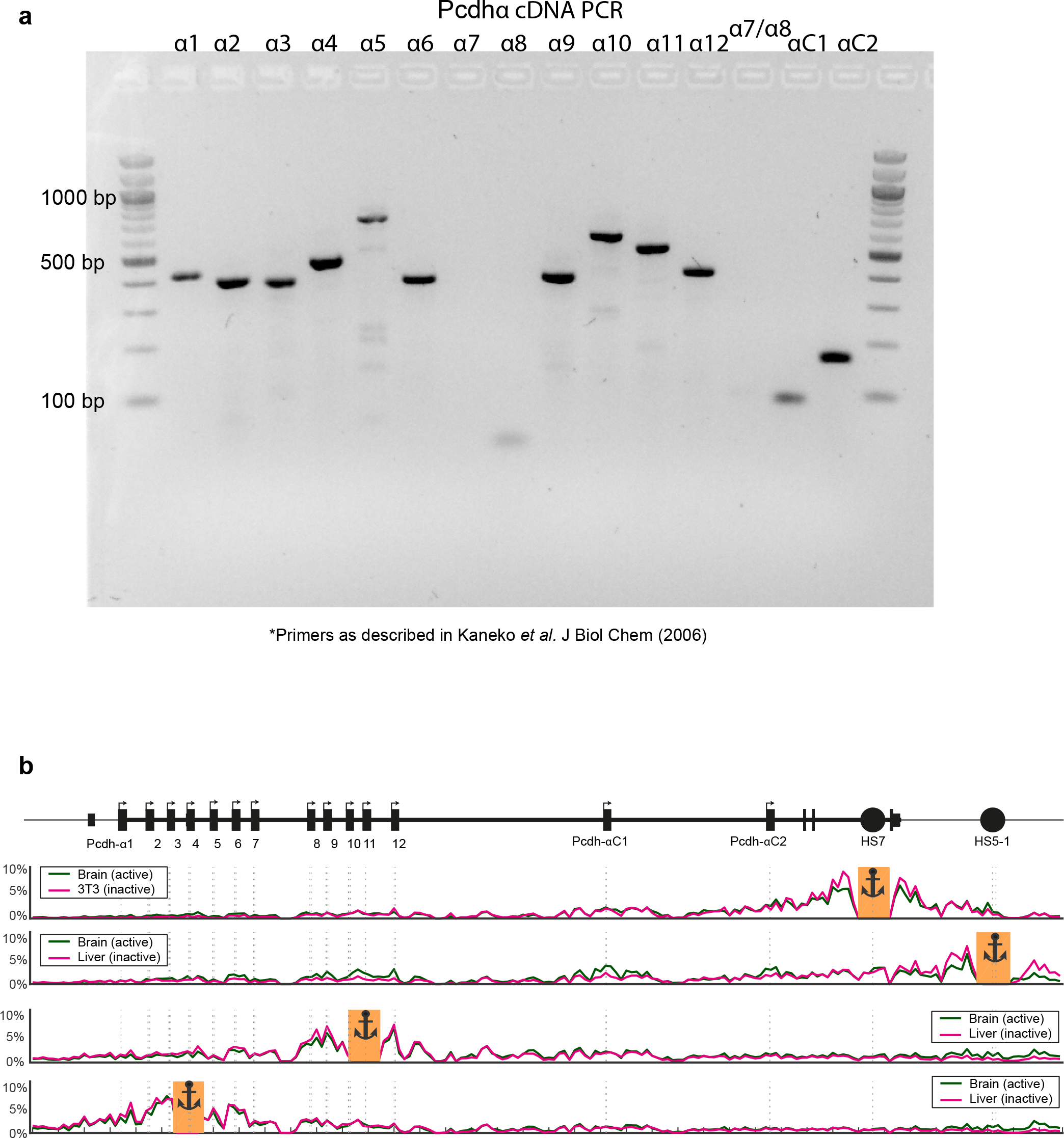
Pcdhα expression in E14.5 fetal brain cells. **a.** Alternative exon-specific primers were used for PCR on cDNA to test which promoters are active in El4.5 fetal brain cells. Primers are listed in Supplementary table 2. **b.** Overall profiles of Pcdh-α4, Pcdh-α11, HS5-1 and HS7 viewpoints in liver (inactive) and brain (active) cells.

**Supplementary Figure 10.**
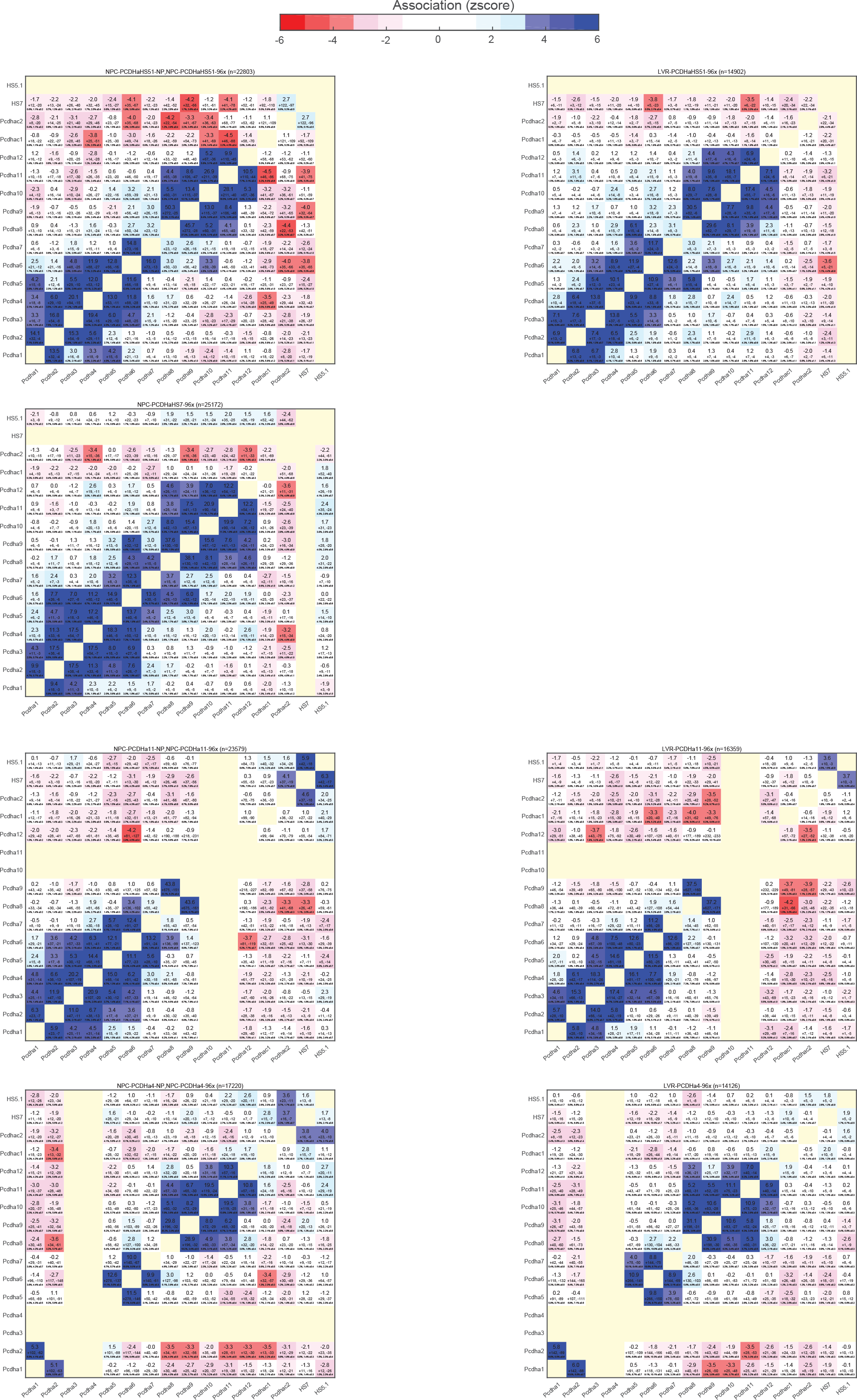
Allelic co-occur frequencies at the active and inactive Pcdhα locus. Complete overview of co-occur significance scores found between any pair of genes and/or enhancers constituents in experiments using Pcdh-α4, α11, HS7 and HS5-1 as viewpoints, in E14.5 fetal liver (Pcdha inactive) and E14.5 fetal brain cells (Pcdhα active).

**Supplementary Figure 11.**
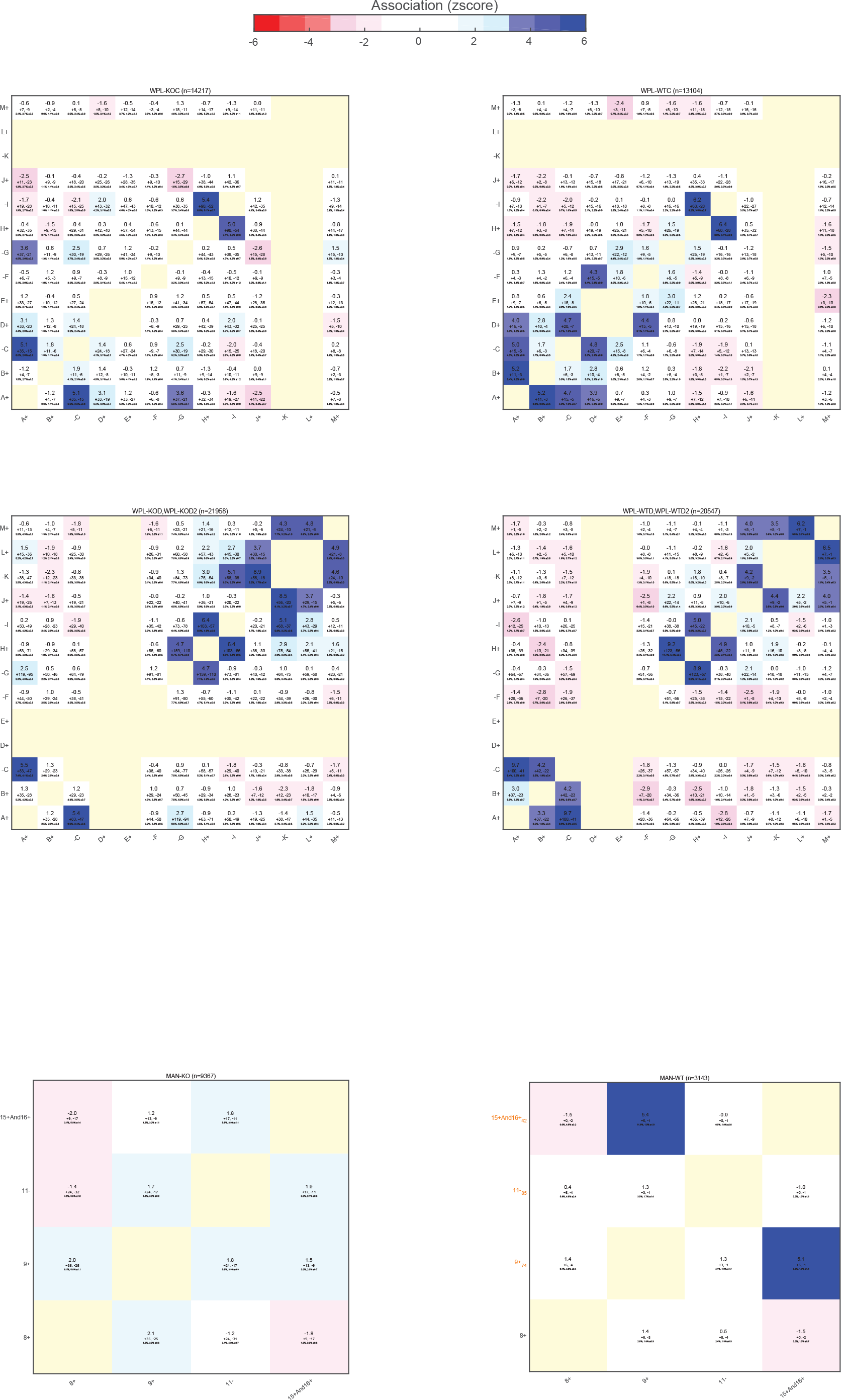
Micro-topologies uncovered in the MAN1A locus in ΔWAPL Hap1 cells. **a.** HiC data obtained in WT and ΔWAPL Hap1 cells, in the MAN 1A locus, showing multiple novel long-range loops formed exclusively in absence of WAPL. Forward and reverse CTCF sites are indicated, as well as the viewpoint used in MC-4C experiments and the CTCF sites used as SOI **b.** Viewpoint-SOI profiles for the MAN 1A viewpoint, using three different CTCF sites as SOI, showing CTCF clustering. z-scores are plotted below the profiles.

**Supplementary Figure 12.**
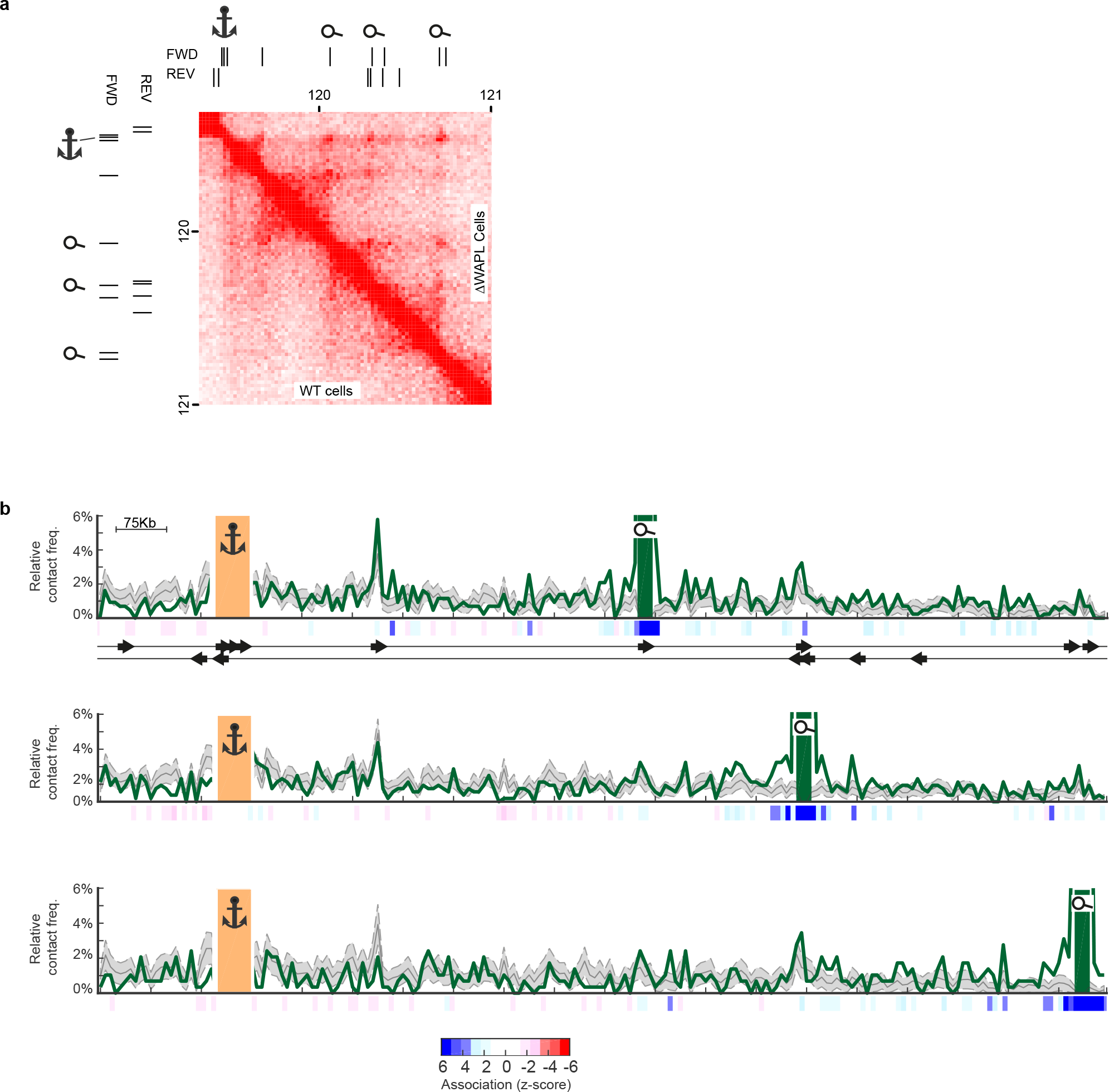
Allelic co-occur frequencies of architectural loops in presence or absence of WAPL. Complete overview of co-occur significance scores found between any pair of CTCF sites in the selected region on chromosome 8, in WT hapl cells and ΔWAPL Hapl cells for both viewpoints (E and K)

### Supplementary Tables

**Supplementary Table 1. Statistics of the MC-4C experiments.** Shown, per experiment, are the total number of raw reads sequenced per experiment, final number of filtered reads (i.e. number of independent alleles analyzed), percentage of total between parentheses, the total number of analyzed contacts and the number of MinION sequencing runs that were pooled for each experiment.

**Supplementary Table 2. Primers used in this study**.

The primers used for each individual viewpoint, FW and RV primers are used for MC-4C PCR, A and B refer to the FW primer used to synthesize the gRNA for the upstream and downstream neighbouring fragment (respectively) of the viewpoint, VP is the primer used to synthesise the gRNA designed on the viewpoint fragment.

### Supplementary method MC-4C computational pipeline

In order reveal the multi-way DNA interactions captured by MC-4C, sequenced reads need to undergo few pre-processing steps. These steps are geared toward validation of read integrity and more crucially eradication of PCR duplicates. The latter, ensures that each utilized read in the downstream analysis is assembled from contacts formed in an individual cell (i.e. unique molecule).

#### 1. Read validity check

To validate fidelity of the sequenced reads, we identified primers as well as their orientations in each respective read. To this end, Bowtie2 v2.2.6 [1] was employed in local alignment mode (settings: -20 -R 3 -N 0 -L 15 -i S,1,0.50 --rdg 2,1 --rfg 2,1 --mp 3,2 --ma 2 -a). We allowed 20% mismatches to take into account the probable errors in nanopore sequencing.

Analysing primer arrangements in sequenced reads showed that some reads are formed by ligation of two individual circles. This observation fostered a correction procedure in which read-ligation events are identified and consequently affected reads were cleaved into two sub-reads. To reduce misidentification of read-ligations events, we allowed no extra sequences between two primers. Additionally, we discarded any reads that contained more than four primers or more than one read-ligation event. These requirements ensured that only those configurations that can be safely rectified go through correction procedure. The produced sub-reads were discarded if their primer configuration did not validate. In this stage, we also discarded any reads that were smaller than 500bp as they may not have enough complexity (i.e. # fragments) to be informative in multi-contact analysis (see **Supplementary table 1** for list of read numbers per experiment).

#### 2. Splitting reads into fragments based on restriction enzyme sequences

A standard aligner (e.g. BWA or Bowtie) capable of split-mapping (i.e. splitting a single query and mapping to multiple coordinates) can be used to map enclosed fragments in each read. To facilitate this, we pre-split the reads into prospective fragments using restriction enzyme sequences. This addition showed improved efficacy in mapping fragments. Due to read errors induced by nanopore sequencing, extra restriction sites (i.e. observing GATC while it should have been GAAC) might be detected. In such cases, the splitted fragments will map alongside each other and further fused together in the merging step of the pipeline (see section 3). On the other hand, a restriction site might be missed. In this case, we relied on the aligner with split-mapping to correctly identify sub-fragments.

#### 3. Mapping reads to reference genome

In order to map produced fragments to reference genome, we utilized BWA-SW v0.7.16a [2] in SW mode (setting: -b 5 -q 2 -r 1 -T 15). Furthermore, the Z-best heuristic of this aligner is set to 10 (i.e. -z 10). This heuristic increases accuracy of the aligner at the cost of speed.

#### 4. Fragment extension and neighbor fusion

Fragments are extended to nearest restriction site (either the 4-cutter or 6-cutter restriction site) in the reference genome. Extension is continued to next restriction site in reference genome, if given fragment has mapped more than 10 bases after identified restriction site. Next, any neighbor fragments in reads that map closer than 30bp in the reference genome are fused together and considered as a single fragment in the rest of analysis. Finally, any fragment with its mapping quality below 20 is considered as unmapped.

#### 5. Duplicate removal (pairwise alignment)

In order to detect PCR duplicates, we utilized a conservative approach which is based on the premise that in MC-4C, fragments far away from viewpoint are unlikely to be found more than once. Therefore, these far-cis/trans fragments can be directly used as Unique Molecular Identifiers (UMI)s [3]. Therefore, if these UMI fragments are identified in two reads, those reads are most likely a duplicate. Schematic of this approach is depicted in Fig.1.

**Fig. 1.**
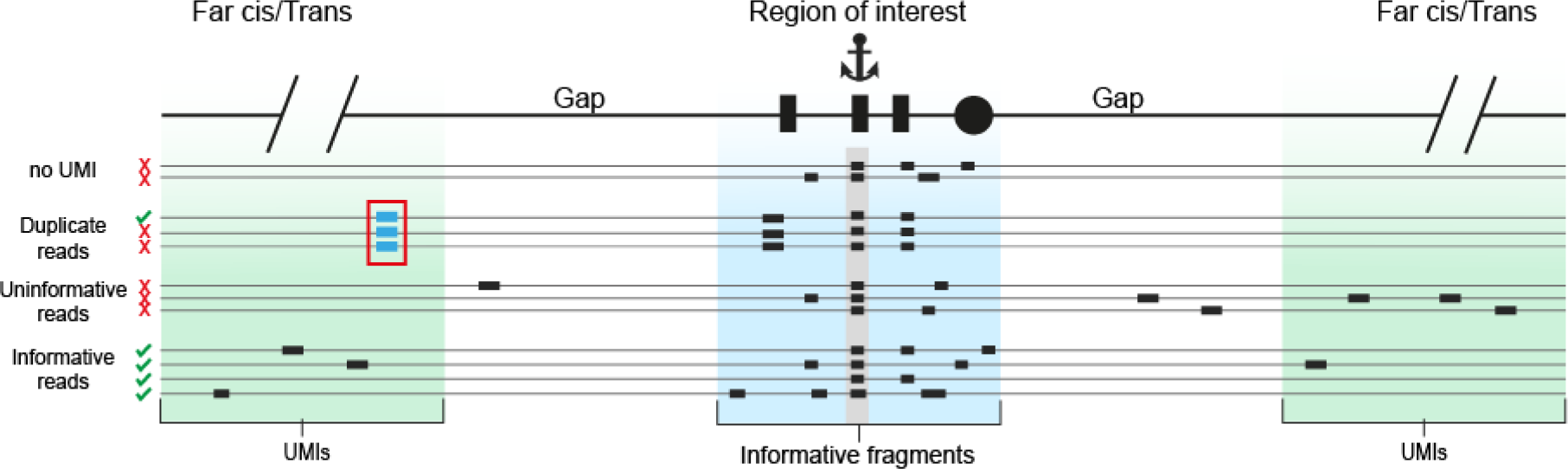
*A schematic representation of PCR filter utilized in MC-4C. Red box represents two reads with identical far-cis/trans fragments. These two reads are considered as duplicates*.

Once a duplicate is found, we removed the read with smaller number of local fragments (i.e. fragments that are mapped within the locus of interest). Finally, reads that have less than two fragments within locus of interest are discarded as they are not informative in multi-way association analysis.

#### 7. Association analysis

We uncovered contact propensity between Viewpoint (V) and two other regions (X and Y) by comparing profiles of circles that contain V and X (Fig.2.a) versus profile of circles that contain V and but not X (Fig.2.b).

**Fig. 2.**
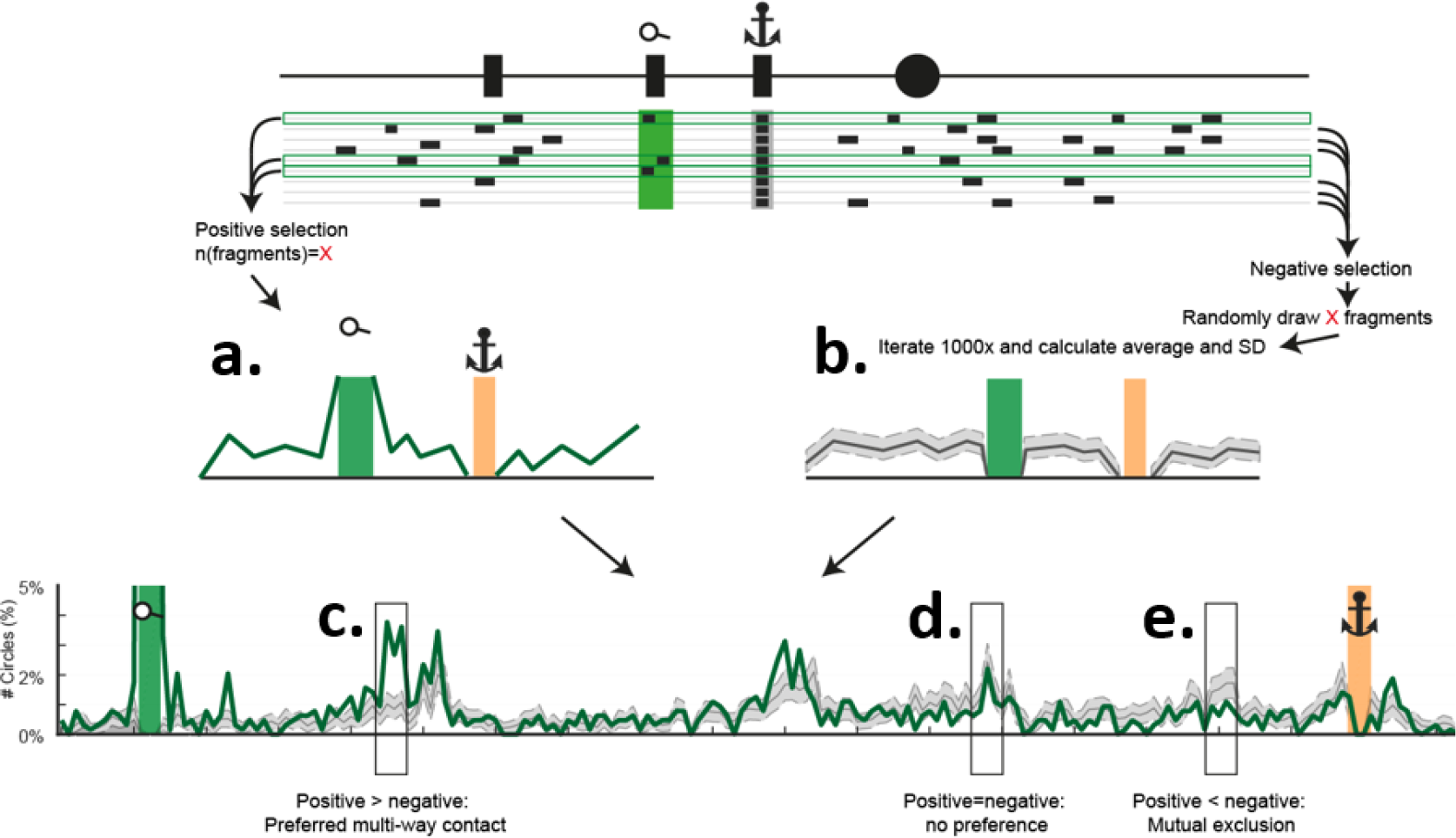
*Schematic overview of association analysis performed for unraveling preferential contacts formed in region of interest*.

We argue that, if there is preferential contact between X and Y (with presence of V), this inclination should be absent when X is not present in the concatemer. Therefore, empowered with multi-way nature of contacts provided by MC-4C, we can directly assess associations between contacts and comparison of contacts with external pairwise data (e.g. Hi-C) [4] is not required.

